# Activation of retinoic acid receptor reduces metastatic prostate cancer bone lesions through blocking endothelial-to-osteoblast transition

**DOI:** 10.1101/2021.12.22.473739

**Authors:** Guoyu Yu, Paul G. Corn, Pengfei Shen, Jian H. Song, Yu-Chen Lee, Song-Chang Lin, Jing Pan, Sandeep K. Agarwal, Theocharis Panaretakis, Maurizio Pacifici, Christopher J. Logothetis, Li-Yuan Yu-Lee, Sue-Hwa Lin

**Affiliations:** Department of Translational Molecular Pathology, The University of Texas M. D. Anderson Cancer Center; Houston, Texas 77030.; Department of Genitourinary Medical Oncology, The University of Texas M. D. Anderson Cancer Center; Houston, Texas 77030.; Translational Research Program in Pediatric Orthopaedics, The Children’s Hospital of Philadelphia; Philadelphia.; Department of Medicine, Section of Immunology Allergy & Rheumatology, Baylor College of Medicine; Houston, Texas 77030.; The University of Texas Graduate School of Biomedical Sciences at Houston; Houston, Texas.

**Author notes:** Co-Corresponding authors: Dr. Sue-Hwa Lin, Department of Translational Molecular Pathology, The University of Texas M. D. Anderson Cancer Center, 1515 Holcombe Blvd., Houston, TX 77030., Dr. Li-yuan Yu-Lee, Department of Medicine, Section of Immunology Allergy & Rheumatology, Baylor College of Medicine, One Baylor Plaza, Houston, TX 77030. These authors contributed equally to this work (Guoyu Yu, Paul G. Corn).

**Keywords:** Bone metastasis, prostate cancer, tumor-induced bone, EC-to-OSB conversion, retinoic acid receptor agonists

## Abstract

Metastatic prostate cancer (PCa) in bone induces bone-forming lesions that contribute to progression and therapy resistance. Currently strategies targeting PCa-induced bone formation are lacking. We previously showed that PCa-induced bone originates from endothelial cells (EC) that have undergone endothelial-to-osteoblast (EC-to-OSB) transition in response to tumor-secreted BMP4. Here, we show that activation of retinoic acid receptor (RAR) inhibits EC-to-OSB transition and reduces PCa-induced bone formation. We found that palovarotene, a RARγ agonist being tested for heterotopic ossification in fibrodysplasia ossificans progressiva, inhibited EC-to-OSB transition and osteoblast mineralization *in vitro*, and decreased tumor-induced bone formation and tumor growth in several osteogenic PCa models. RARα/β/γ isoform knockdown in 2H11 ECs blocked EC-to-OSB transition and osteoblast mineralization. Pan-RAR agonist ATRA inhibited MycCaP-BMP4-induced bone formation and tumor growth under castration. Furthermore, palovarotene or ATRA reduced plasma Tenascin C, a factor secreted by EC-OSB cells, which may be used to monitor treatment response. Mechanistically, BMP4-activated pSmad1 forms a complex with RAR in the nucleus of 2H11 cells. RAR activation by palovarotene or ATRA causes pSmad1 degradation by recruiting E3-ubiquitin ligase Smurf1 into the nuclear pSmad1/RARγ complex. Our findings suggest that palovarotene can be repurposed to target PCa-induced bone formation to improve clinical outcomes for bone metastasis.

## INTRODUCTION

Metastatic progression in bone signifies the lethal progression of advanced prostate cancer (PCa). Currently approved therapeutic agents for PCa bone metastasis only modestly affect patient survival (1). Thus, there is an urgent need to develop strategies that improve therapy outcomes. PCa bone metastasis is frequently associated with osteoblastic bone-forming lesions (2), indicative of cross-talk between metastatic tumor cells and the stromal component within the tumor microenvironment. Lin et al. (3) showed that reduction of tumor-induced bone formation resulted in decreased osteogenic tumor growth in a genetic mouse model. Inhibition of tumor-induced bone formation, by the BMP receptor inhibitor LDN-193189 (4) or an anti-BMP6 antibody (5), reduced tumor growth in mouse models. These observations support a role of osteoblasts in PCa progression in bone. In addition, PCa-induced bone formation increases therapy resistance (6). Thus, inhibition of PCa-induced bone formation may improve therapy outcomes.

Therapies targeting PCa-induced bone formation are limited. Bisphosphonates and RANK-L inhibitors reduce skeletal-related events but have failed to improve overall survival (7). Several bone-targeting radionuclides, including Sr89 and Samarium-153 EDTMP, were initially developed as bone-targeted therapy for PCa bone metastasis, but elicited significant bone marrow suppression that limited their further clinical development (8). In contrast, Radium-223, a bone targeting alpha emitter that remodels bone by inducing cell death in osteoblasts, prolongs survival of patients with bone metastasis (9). The life prolonging effect of Radium-223, while modest, is consistent with the hypothesis that PCa-induced bone formation plays a role in prostate tumor progression in bone. Based on this, additional strategies to reduce tumor-induced bone formation may further improve therapy outcome for PCa bone metastasis.

Until recently, the mechanism of PCa-induced aberrant bone formation was thought to result from the expansion of existing osteoblasts. However, we recently reported an alternative mechanism whereby PCa-induced aberrant bone formation occurs via endothelial-to-osteoblast (EC-to-OSB) transition mediated by tumor-secreted BMP4 (3). Inhibition of EC-to-OSB transition through endothelial-specific deletion of osterix, an osteoblast-specific transcription factor, reduced PCa-induced bone formation and tumor growth (3). Targeting this unique EC-to-OSB transition may reduce PCa-induced bone formation.

This new understanding of EC-to-OSB transition in PCa–induced bone formation has revealed similarities in the mechanism(s) of PCa–induced bone and ectopic bone arising from the genetic disease fibrodysplasia ossificans progressiva (FOP), also known as stoneman disease (10, 11). FOP is a rare genetic disorder in which muscle and connective tissues are gradually replaced by bone, forming bone outside the skeleton (heterotopic ossification) that constrains movement and is debilitating. FOP is caused by a hereditary gain-of-function genetic mutation (R206H) in the BMP type I receptor ALK2 that activates BMP signaling pathways, leading to the formation of ectopic bone (12),(13). Mechanistically, the cell source for ectopic bone formation was shown to be ECs that undergo endothelial-to-mesenchymal transition (EndoMT) to become mesenchymal stem cells, which further differentiate into bone tissues (10). Thus, in both diseases the aberrant bone originates from ECs with over-activated BMP signaling. The similarities in the mechanism have led to our hypothesis that therapies to prevent heterotopic ossification in FOP could be repurposed for inhibiting PCa–induced aberrant bone formation.

Various strategies have been developed for the treatment of FOP. Most of them target the unique changes caused by ALK2 mutation (R206H) (14). Recently, studies by Shimono et al.(15) showed that compounds that specifically activate the nuclear receptor, retinoic acid receptor gamma (RARγ), are effective in inhibiting chondrogenesis and preventing heterotopic ossification in FOP. As cartilage development normally requires an un-liganded RARγ repressor function,(16) the presence of ligands (RARγ agonists) may interfere with transcriptional programming for chondrogenesis and osteogenesis (16). Currently, palovarotene (Palo) is being evaluated as therapy for FOP patients in a phase III clinical trial (NCT03312634). Palo may be repurposed to inhibit PCa–induced aberrant bone formation.

RARγ is a nuclear hormone receptor that binds to specific DNA response elements to regulate gene transcription. In the absence of ligands, RARγ acts as a transcriptional repressor by recruiting co-repressors to target promoters. When ligands are present, RARγ interacts with coactivators to activate gene transcription (17, 18). RARγ is known to regulate major developmental processes (19). Retinoic signaling plays a critical role in modulating skeletal development (20–22). Overexpression of RARα in transgenic animals leads to appendicular skeletal defects (23). A loss of retinoid receptor–mediated signaling is necessary and sufficient for expression of bone phenotype (24). Thus, retinoic acid signaling inhibits skeletal progenitor differentiation (22).

In this study, we examined whether Palo may be repurposed to inhibit PCa–induced aberrant bone formation. We found that RAR agonists, including Palo and all-trans-retinoic acid (ATRA) inhibit BMP4-induced EC-to-OSB transition and bone formation. In addition, RARs regulate BMP4-induced EC-to-OSB transition through a non-canonical RAR mechanism that is independent of RAR’s transcription factor function. Further, we found that plasma levels of Tenascin C, a factor secreted by EC-OSB cells, can be used for monitoring treatment response. ATRA treatment also does not lead to unwanted bone loss in healthy non-tumor-bearing bones. Thus, Palo and ATRA represent new agents for the treatment of osteoblastic bone metastasis of PCa.

## RESULTS

### RARγ agonist palovarotene inhibits EC-to-OSB transition

2H11 endothelial cells can be induced to undergo endothelial-to-osteoblast (EC-to-OSB) transition by treating with BMP4 (100 ng/mL) for 48 h, based on the expression of markers OSX, an osteoblast cell fate determination factor, and osteocalcin (Fig. 1A, B). The addition of palovarotene (Palo) significantly inhibited BMP4-induced EC-to-OSB transition as shown by the inhibition of OSX and osteocalcin expression at both mRNA and protein levels (Fig. 1A, B). Furthermore, Palo treatment led to the inhibition of mineralization, as indicated by Alizarin Red (Fig. 1C) and von Kossa staining (Fig. 1D), compared to BMP4 treatment alone when cultured in osteoblast differentiation medium for 21 days. Palo alone has no effect on EC-OSB transition. These results suggest that Palo inhibits BMP4-induced EC-to-OSB transition.

**Figure 1.**
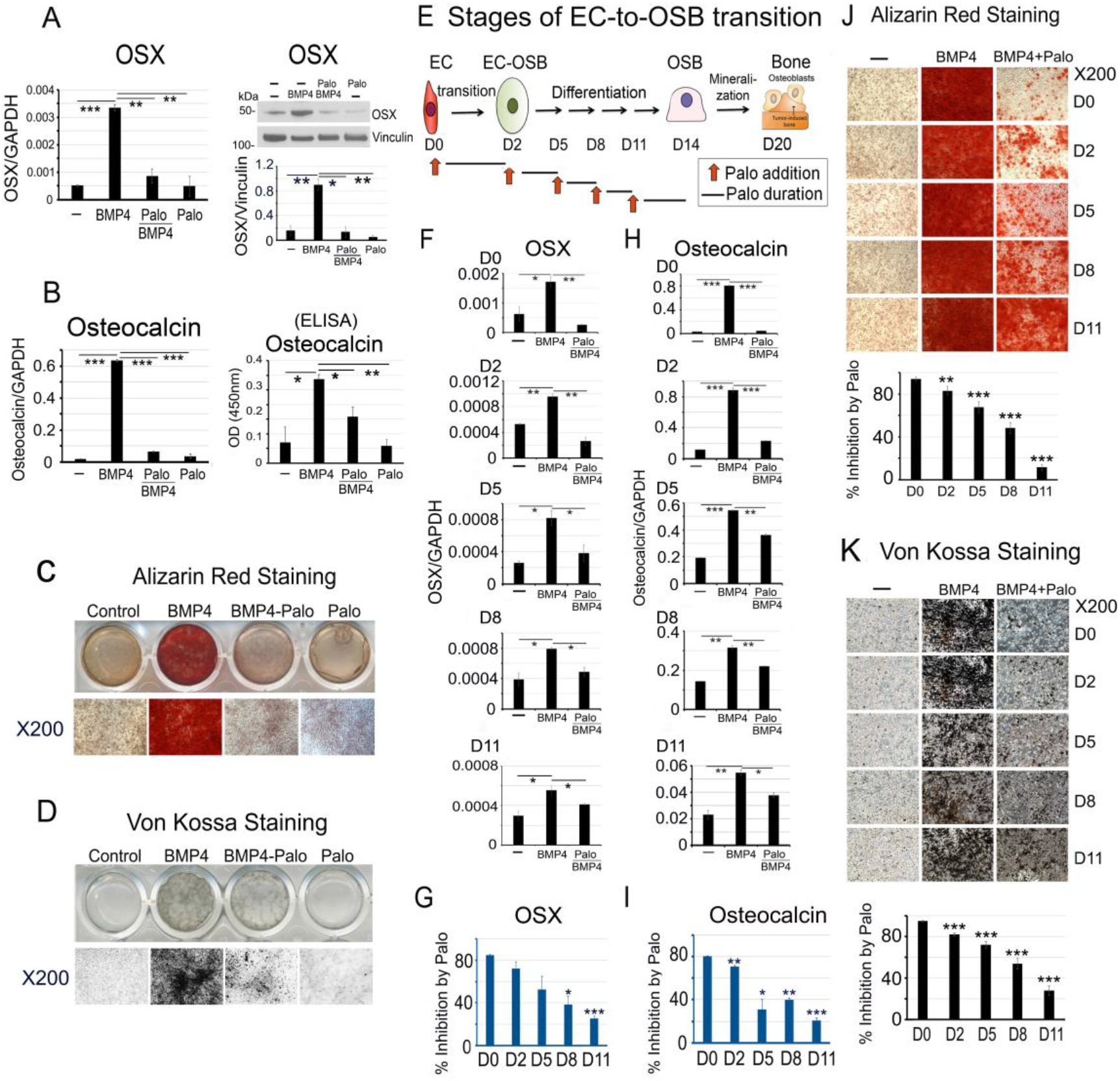
Palovarotene inhibits BMP4-induced EC-to-OSB transition in 2H11 cells. (A-D) 2H11 endothelial cells were treated with BMP4 (100 ng/mL) to induce EC-to-OSB transition. Palovarotene (Palo, 1 μM) treatment effects were measured on (A) OSX mRNA by qRT-PCR and protein by western blot, (B) osteocalcin mRNA by qRT-PCR and protein in conditioned medium by ELISA, (C) mineralization by Alizarin Red staining, and (D) mineralization by Von Kossa staining. (E-K) Palo effects at different stages of EC-to-OSB transition. (E) Phases of EC-to-OSB transition in 2H11 cells following BMP4 treatment. Red arrows indicate Palo addition at different times followed by incubation for 3 days. (F) OSX mRNA expression when Palo was added at D0, D2, D5, D8 or D11. (G) Percent inhibition of OSX expression by Palo in cells in (F) when compared to BMP4-treated only. Percent inhibition was calculated as 100-[(BMP4+Palo)/BMP4]x100. (H) Osteocalcin mRNA expression in cells treated as in (F). (I) Percent inhibition of osteocalcin expression by Palo determined as in (G). (J) Palo on mineralization when added at D0, D2, D5, D8 or D11, as determined by Alizarin Red staining or (K) Von Kossa staining. Staining was quantified using ImageJ. *p<0.05, **p<0.01, ***p<0.001 by Students *t*-test.

### Inhibition by palovarotene occurs in the early stages of EC-to-OSB transition

2H11 cells undergo EC-to-OSB transition in several stages. Upon BMP4 treatment, 2H11 cells transition into OSB by day 2 (days 0-2, Transition Phase) as determined by OSX and osteocalcin expression. EC-OSB hybrid cells then undergo OSB differentiation and express bone matrix proteins (days 3-14, Differentiation Phase), and finally mineralize by day 20 (Mineralization Phase) (Fig. 1E). To examine whether Palo could inhibit BMP4-induced bone formation after EC-to-OSB transition has already been initiated, Palo was added to 2H11 cells at different stages of EC-to-OSB transition. Palo was most effective during the early Transition Phase (D0 – D2), reaching about 80 – 90% inhibition of BMP4-stimulated activities (Fig. 1G, 1I). Palo still inhibited OSX mRNA (Fig. 1F) and osteocalcin mRNA (Fig. 1H), albeit with decreasing efficacy, when added to the cultures on D2, D5, D8, or D11 (Fig. 1F-1I), by which time inhibition was reduced to about 20% that of BMP4-treated cells. Similarly, inhibition of mineralization was also most effective during early transition and less effective when Palo was added in later phases of EC-to-OSB transition (Fig. 1J and 1K). These observations suggest that Palo is most effective in inhibiting early stages of BMP4-induced EC-to-OSB transition.

### Palovarotene reduces PCa-induced aberrant bone formation in xenograft animal models

We employed three osteogenic xenograft models to examine the effects of Palo on PCa-induced bone formation. In the first model, C4-2b-BMP4 cells, which form osteogenic tumors (3), were injected subcutaneously into SCID mice (Fig. 2A). Mice were treated with or without Palo (2 mg/kg/day) by oral gavage. Palo treatment significantly reduced tumor-induced bone formation compared to those in vehicle-treated control tumors, as determined by Goldner’s Trichrome staining (p=0.002) (Fig. 2B). We also observed that Palo led to a moderate decrease in tumor sizes compared with controls, as determined by bioluminescence imaging (Fig. 2C) and tumor weight (Fig. 2D). However, Palo did not have a significant effect on C4-2b-BMP4 cell proliferation in vitro (Supplementary Fig. 1A) or on mouse body weight (Supplementary Fig. 1B). These observations suggest that Palo reduced tumor-induced bone formation and moderately reduced tumor size, consistent with previous observation that PCa-induced bone supports prostate tumor growth (4, 5).

**Figure 2.**
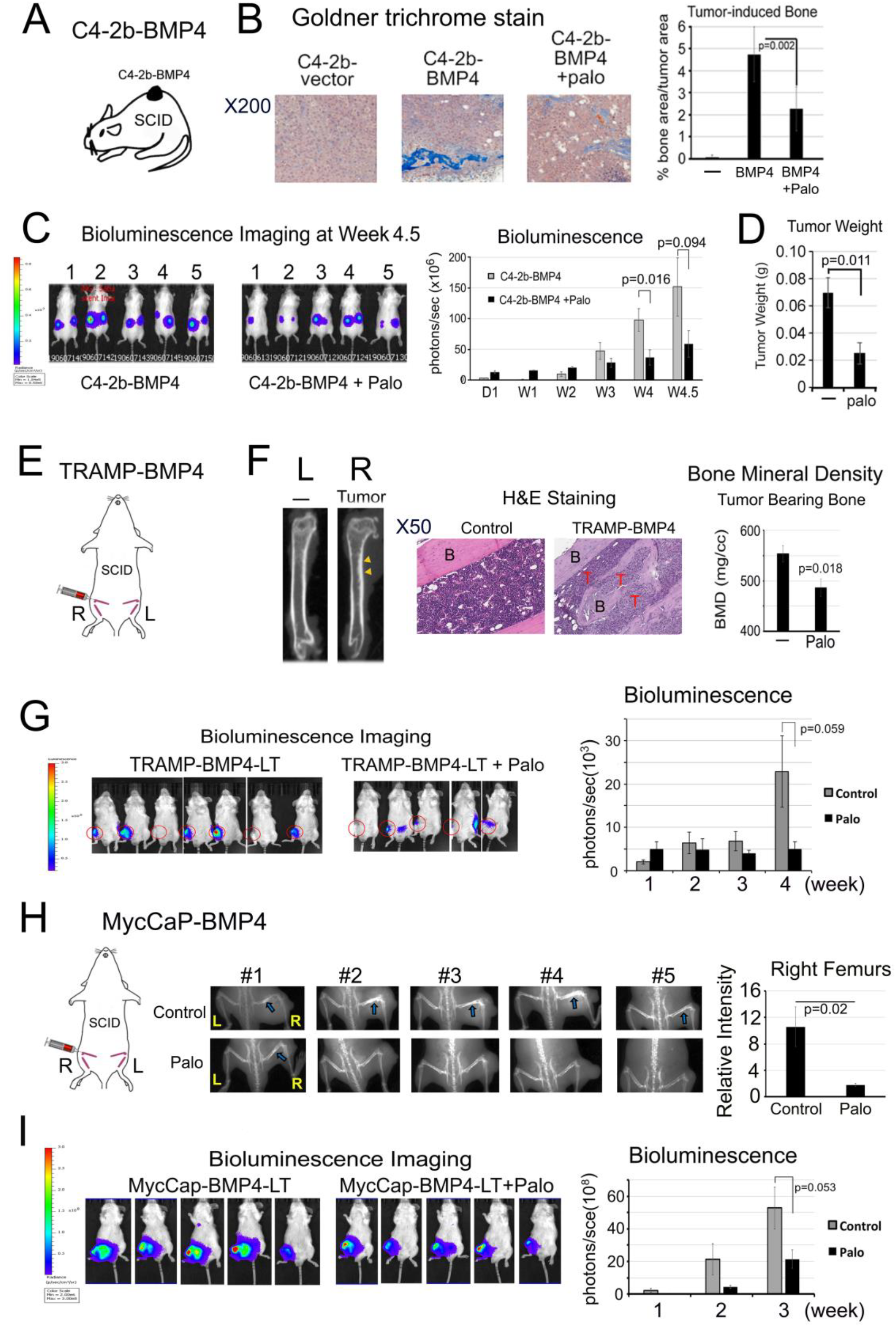
Palovarotene treatment reduces PCa-induced bone formation and tumor growth in SCID mice. (A) C4-2b-BMP4 subcutaneous implantation model. (B) Goldner Trichrome staining of C4-2b-BMP4 tumors and quantification using ImageJ. (C) Tumor size measurement by BLI in mice with C4-2b-BMP4 tumors. (D) Tumor weight at the termination of the study. control (n=5), Palo-treated (n=5). (E) TRAMP-BMP4 intrabone injection model. (F) MicroCT of femurs with or without TRAMP-BMP4 tumors (left). Histology of TRAMP-BMP4-induced bone compared with the non-tumor bearing bone (middle). Palo effect on bone mineral density of tumor-bearing femurs (right). B(bone), T(tumor). (G) TRAMP-BMP4 tumor size measurement by BLI. Control (n=7); Palo-treated (n=5). (H) MycCaP-BMP4 intrabone injection model (left). X-Ray analysis of osteoblastic bone response induced by MycCaP-BMP4 in femurs with or without Palo on (middle) and relative intensity quantified by ImageJ (right). Blue arrows, tumor bearing bone in the control group. (I) MycCaP-BMP4 tumor size measurement by BLI. Control (n=5); Palo-treated (n=5).

In a second model, we injected osteogenic TRAMP-BMP4 cells intrafemorally into SCID mice and treated these tumor-bearing mice with or without Palo (Fig. 2E). TRAMP-BMP4 cells were generated by transfecting TRAMP-C2 cells (25) with cDNA encoding mouse BMP4 and express high levels of BMP4 mRNA and protein compared with control TRAMP-C2 cells (Supplementary Fig. 1C). Micro-CT analysis showed that TRAMP-BMP4 increased the bone mineral density of the tumor-bearing femurs compared to the control un-injected legs (Fig. 2F, left) and histology showed TRAMP-BMP4 induced bone formation in the bone marrow (Fig. 2F, middle). Palo significantly reduced the bone mineral density, as determined by μCT, in the tumor-bearing femurs compared with those in non-treated mice (Fig. 2F, right) and moderately decreased PCa growth in bone at week 4 by bioluminescence, although the decrease did not reach statistical significance (Fig. 2G). Palo did not have a significant effect on TRAMP-BMP4 cell proliferation in vitro (Supplementary Fig. 1D). In this experiment, Palo led to a small decrease in mouse body weight compared to untreated mice (Supplementary Fig. 1E). These results suggest that Palo treatment reduces TRAMP-BMP4-induced bone formation in mouse femurs.

In a third model, MycCaP-BMP4 cells were injected intrafemorally into SCID mice and treated with or without Palo (Fig. 2H). MycCaP cells, derived from prostate tumor of Hi-Myc mice (26), do not express detectable BMP4. Upon transfection with mouse BMP4 cDNA, MycCaP-BMP4 cells expressed a high level of BMP4 proteins relative to MycCaP cells (Supplementary Fig. 1F). X-Ray analysis showed that MycCaP-BMP4 in femurs induced an osteoblastic bone response in tumor-bearing femurs, and Palo significantly reduced this response (Fig. 2H). Palo also exhibited a trend towards reduced tumor growth by bioluminescence (Fig. 2I). Palo treatment did not have a significant effect on MycCaP-BMP4 cell proliferation in vitro (Supplementary Fig. 1G) and mouse body weight (Supplementary Fig. 1H). Together, the combined results from these three osteogenic xenograft models suggest that Palo treatment reduces PCa-induced bone formation accompanied with a trend of decreased tumor growth.

### Palovarotene reduces BMP4-stimulated pSmad1 levels in pSmad1/RARγ complex

Smad1 phosphorylation is one of the downstream BMP4 signaling pathways in 2H11 cells (27). pSmad1 levels were significantly increased by BMP4, and Palo treatment reduced BMP4-stimulated pSmad1 levels (Fig. 3A). Total Smad1 was also increased by BMP4 and slightly decreased by Palo (Fig. 3A). BMP4 also increased nuclear localization of pSmad1, which was decreased by Palo (Fig. 3B). Similar patterns of distribution were observed with Smad1 (Fig. 3B). In contrast, total and nuclear RARγ levels were not affected by BMP4 or Palo (Fig. 3A, B). These observations suggest that Palo treatment leads to a decrease in BMP4-stimuated nuclear pSmad1 levels.

**Figure 3.**
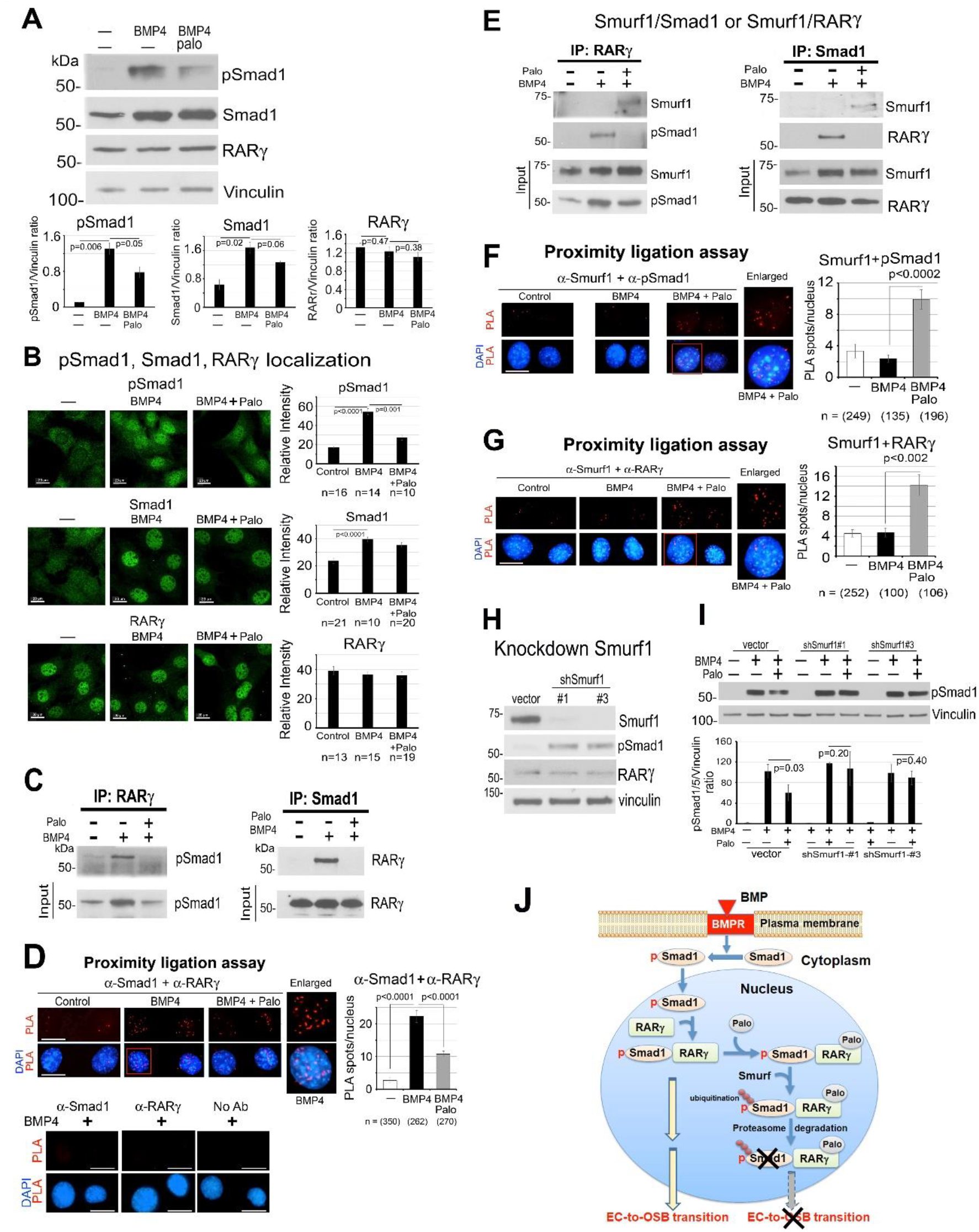
Palovarotene decreases BMP4-stimulated pSmad1. (A) pSmad1, Smad1 and RARγ protein levels in 2H11 cells treated as indicated for 6 h. Vinculin, loading control. Blots were quantified by ImageJ. (B) Immunofluorescence of pSmad1, Smad1, and RARγ in nucleus of cells in (A). Relative intensity was quantified by ImageJ. n, number of nuclei examined. All bars, 20 μm. (C) Co-immunoprecipitation of RARγ and pSmad1 from nuclear extracts. (D) Proximity ligation assay for the interaction of RARγ and pSmad1. Red spots, PLA signals in the nucleus using anti-Smad1 and anti-RARγ antibodies. Single or no antibodies were used as controls. (E) Smurf1 in co-immunoprecipitations with RARγ or pSmad1 using nuclear extracts. (F-G) Proximity ligation assay for the interaction of Smurf1 with pSmad1 (F) or RARγ (G). (H) Smurf1 knockdown in 2H11-shSmurf1#1 or #3 clones. (I) pSmad1 levels in 2H11-shSmurf1 knockdown clones. (J) Graphical summary of mechanism of Palo-mediated inhibition of EC-to-OSB transition. BMP4 treatment leads to an increase in pSmad1 that translocates into the nucleus, where pSmad1 and RARγ form a complex. Palo treatment results in the recruitment of E3-ubiquitin ligase Smurf1 to the pSmad1/RARγ complex, leading to pSmad1 degradation and inhibition of EC-to-OSB transition.

Using nuclear extracts, we showed that immunoprecipitation of RARγ pulled down pSmad1, while the reciprocal immunoprecipitation of Smad1 pulled down RARγ (Fig. 3C), suggesting that pSmad1 forms a complex with RARγ in BMP4-treated 2H11 cells. Further, *in situ* proximity ligation assay (PLA) for the potential physical association between pSmad1 and RARγ showed a significant increase in PLA signals, visualized as individual fluorescent spots, in the nucleus of BMP4-treated 2H11 cells but not in control cells (Fig. 3D). No PLA signals were observed in BMP4-treated cells incubated with a single antibody alone as controls (Fig. 3D). Palo was found to reduce BMP4-mediated nuclear pSmad1/RARγ complex formation by both co-immunoprecipitation (Fig. 3C) and proximity ligation assays (Fig. 3D). These results suggest that Palo inhibits BMP4 stimulated pSmad1/RARγ nuclear complex formation.

### Palovarotene increases Smurf1-mediated phospho-Smad1 degradation

We postulated that Palo inhibits BMP4 stimulated pSmad1/RARγ nuclear complex formation by enhancing its degradation via Smad ubiquitination regulatory factor1 (Smurf1) (28–30). We tested this by co-immunoprecipitations and found that Palo treatment led to the recruitment of E3-ubiquitin ligase Smurf1 to the nuclear pSmad1/RARγ complex (Fig. 3E). Further, *in situ* PLA assays showed a significant increase in Smurf1 association with pSmad1 (Fig. 3F) as well as Smurf1 association with RARγ (Fig. 3G) in the nucleus of BMP4-treated 2H11 cells, only in the presence of RAR activation by Palo but not BMP4 alone. Together, these observations suggest that Palo recruits Smurf1 to the nuclear RARγ/pSmad1 complex.

We knocked down Smurf1 in 2H11 cells and found an increase in pSmad1 levels in 2H11-shSmurf1#1 and #2 clones compared to vector control (Fig. 3H), consistent with a role of Smurf1 in regulating pSmad1 protein levels. In 2H11-shSmurf1 clones, the Palo-mediated decrease in pSmad1 was abrogated compared with vector controls (Fig.3I), suggesting that Palo mediates pSmad1 degradation through Smurf1, likely through pSmad1 ubiquitination for proteasomal degradation. Together, our studies suggest that RAR activation by Palo causes BMP4-stimulated phospho-Smad1 degradation by recruiting the E3-ubiquitin ligase Smurf1 into the nuclear pSmad1/RARγ complex (Fig. 3J).

### Palovarotene alters the BMP4-induced transcriptome

We characterized the Palo-regulated transcriptome by RNAseq, which showed that mRNA levels of 591 genes were upregulated more than 1.5-fold by BMP4 when compared to 2H11 cells (2H11-BMP4 vs 2H11) (Fig. 4A). By using fold enrichment to select for the top 215 pathways followed with lowest false discovery rate, pathway analysis showed that regulation of osteoblast differentiation, bone morphogenesis and bone mineralization were the most significantly upregulated pathways by BMP4 (Fig. 4B), consistent with ECs transition into OSBs. Upon Palo treatment, mRNA levels of 756 genes were downregulated compared to BMP-treated sample ([BMP4+Palo] vs BMP4) (Fig. 4A) (see GSE168157). Pathway analysis showed that genes involved in regulation of ossification were most significantly downregulated by Palo (Fig. 4C), consistent with an effect of Palo on inhibiting bone formation.

**Figure 4.**
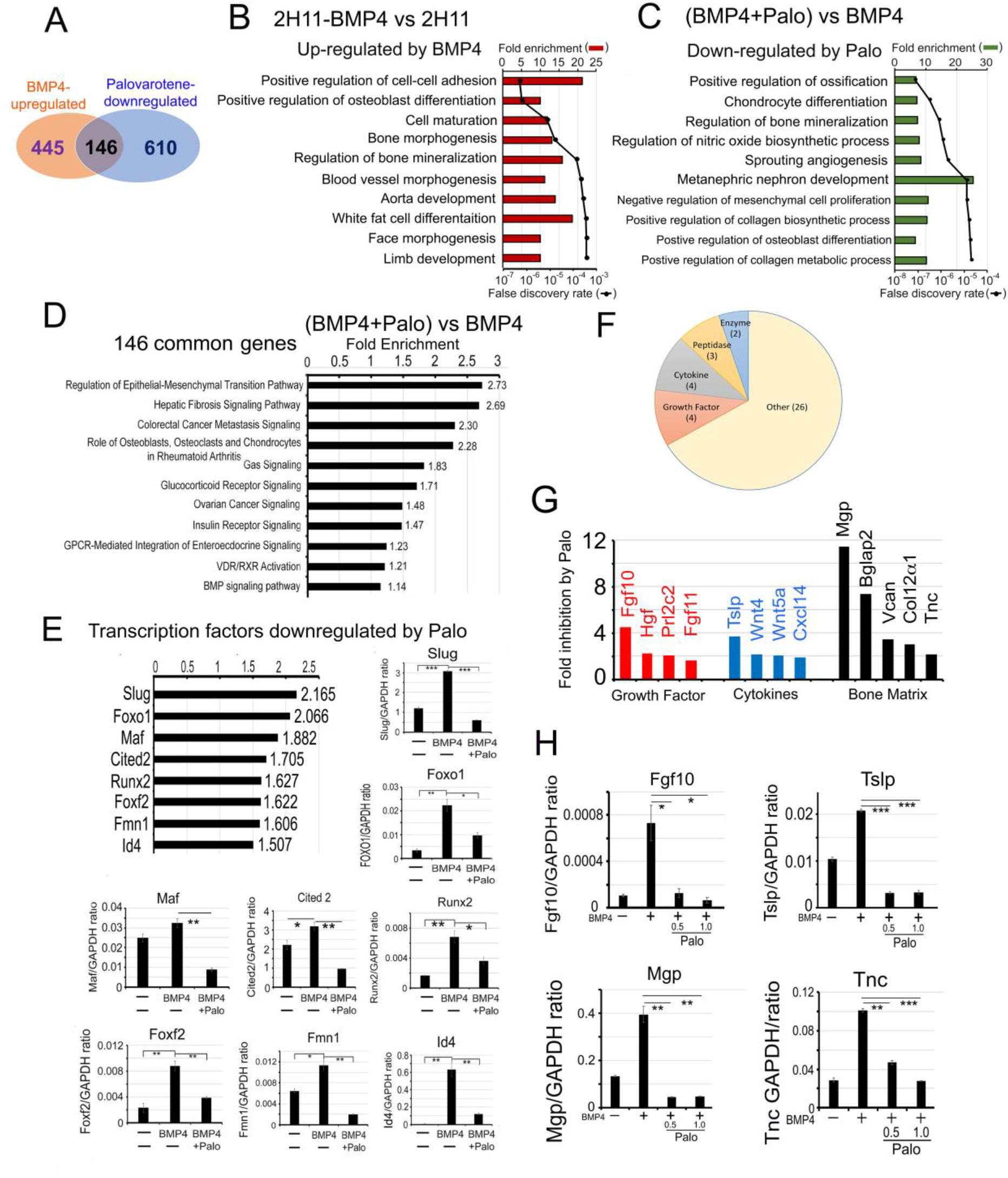
Palovarotene alters the BMP4-induced transcriptome. (A) Venn diagram of number of genes whose levels are upregulated by BMP4 and downregulated by Palo in 2H11 cells. Pathway analyses of genes upregulated by BMP4 (B), downregulated by Palo (C), and up-regulated by BMP4 but down-regulated by Palo (Palo-regulated genes) (D). (E) Transcription factors upregulated by BMP4 and downregulated by Palo. qRT-PCR for mRNAs of eight transcription factors as indicated. (F) Category of BMP4-inducible secreted proteins that are inhibited by Palo. (G) BMP4-stimulated secreted proteins, which we termed “osteocrines” (6), which are decreased by Palo. (H) qRT-PCR for mRNAs of Fgf10, Tslp, Mgp, and Tnc in response to BMP4 and Palo as indicated.

We identified 146 transcripts that were upregulated by BMP4 and downregulated by Palo (Fig. 4A). Pathway analysis showed that these 146 transcripts are enriched in genes involved in regulation of epithelial to mesenchymal transition and the function of osteoblasts, osteoclasts and chondrocytes in Rheumatoid Arthritis (Fig. 4D), consistent with a role of Palo in inhibiting EC-to-OSB transition. Further, we found that 8 transcription factors that play critical roles in cell fate change, including Slug, Foxo1, and Id4, are downregulated by Palo (Fig. 4E). qRT-PCR confirmed that the mRNAs of these transcription factors were upregulated by BMP4 and downregulated by Palo (Fig. 4E). Interestingly, Id4 is a DNA-binding protein that is a downstream effector of BMP signaling and regulates various cellular processes (31). Slug, also known as Snai2, is a cell fate determinant (32). Foxo1 regulates osteoblast numbers, bone mass, and the differentiation of mesenchymal cells into osteoblasts (33). These results suggest that the binding of Palo to RARγ alters Smad1 signaling, leading to transcriptional changes of several cell fate related transcription factors involved in osteoblastogenesis.

We also identified 39 secreted proteins upregulated by BMP4 and downregulated by Palo (Fig. 4F and Supplementary Table 1). These proteins may be involved in stromal-PCa communication critical for PCa progression in bone. These include growth factors, cytokines, proteases, and bone matrix proteins (Fig. 4G and Supplementary Table 1), collectively termed “osteocrines” (6) secreted by EC-OSB cells. The mRNA levels of Fgf10, Tslp, Mgp and Tnc were upregulated by BMP4 and significantly downregulated by Palo in 2H11 cells (Fig. 4H). Fgf10 is required for limb and lung development (34, 35). Tenascin C (TNC), a bone matrix protein, plays a role in inducing stemness and increasing metastatic potential of tumor cells (36). The cytokine TSLP, thymic stromal lymphopoietin, is involved in the maintenance of Th2-type homeostasis. TSLP has recently been found to be involved in the induction and progression of a variety of tumors (37, 38). These secreted factors may be involved in Palo-mediated inhibition of tumor growth and can be used as serum biomarkers for monitoring response to Palo treatment.

### RARα and RARβ are also involved in EC-to-OSB transition

RAR family proteins contain 3 isoforms, RARα, RARβ and RARγ. Palo is a synthetic retinoic acid that has a high affinity for RARγ. It is not clear whether RARα and/or RARβ are involved in EC-to-OSB transition is unclear. We determined that 2H11 cells express all three isoforms by qRT-PCR (Fig. 5A-C). We found that knockdown of either RARα, β, or Rγ (Fig. 5A-C) significantly reduced BMP4-induced osteocalcin expression in the respective 2H11-shRARα, β and γ clones (Fig. 5A-C). Thus, all three RAR isoforms are involved in BMP4-induced EC-to-OSB transition. Knockdown of RAR isoforms also led to an inhibition of BMP4-induced pSmad1 levels (Fig. 5D), suggesting similar mechanisms of inhibition mediated by the three RAR isoforms.

**Figure 5.**
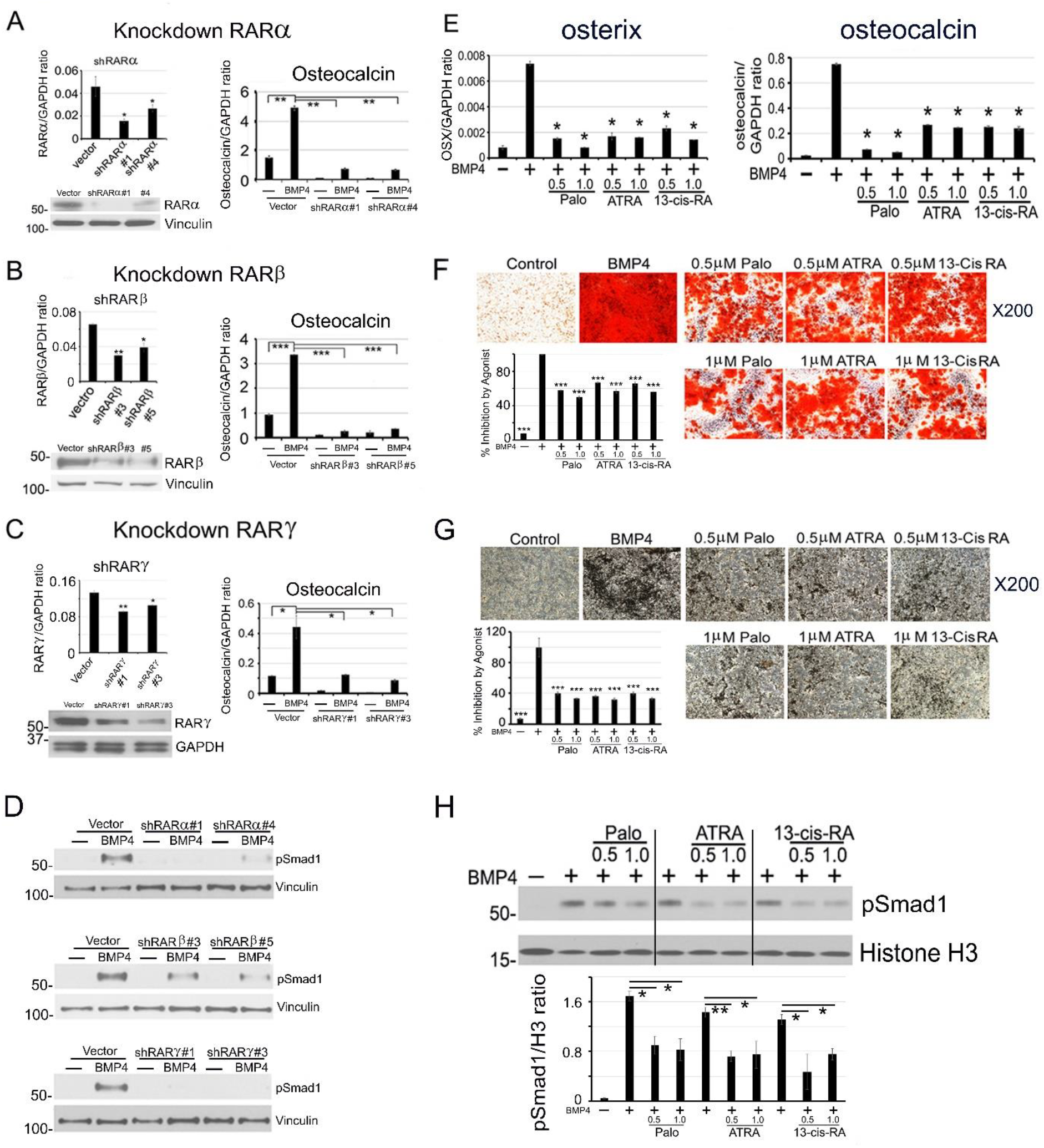
All three RAR isoforms are involved in EC-to-OSB transition. (A-C) RARα, RARβ, or RARγ knockdown in 2H11 cells by shRNA. RAR knockdowns by qRT-PCR and western blot (left). RAR knockdowns on BMP4-stimulated osteocalcin expression (right). (D) RAR knockdowns on BMP4-stimulated nuclear pSmad1 activation. (E-G) ATRA or 13-cis-retinoic acid treatment on BMP4-stimulated osterix and osteocalcin expression (E), mineralization measured by Alizarin red (F), or von Kossa (G) staining. *p<0.05, **p<0.01, ***p<0.001. (H) pSmad1 levels in BMP4-stimulated 2H11 cells treated with Palo, ATRA or 13-cis-RA. Histone H3, nuclear protein loading control. All the samples were run on the same gel.

### ATRA and 13-cis-retinoic acid inhibit EC-to-OSB transition

We next examined if retinoic acids with broad spectrum for the different RAR isoforms can also inhibit BMP4-induced EC-to-OSB transition. We found that tretinoin (ATRA) and isotretinoin (13-cis-RA), which are agonists for all RAR isoforms and are used in the clinic (39), both inhibited EC-to-OSB transition in a dose-dependent manner (Fig. 5E-G), to a similar extent as that of Palo. ATRA and 13-cis-RA also inhibited BMP4-induced nuclear pSmad1 to levels similar as that by Palo (Fig. 5H). These results suggest that retinoic acid agonists, including Palo, ATRA and 13-cis-RA, can inhibit BMP4-induced EC-to-OSB transition.

### ATRA inhibits BMP4-induced bone formation following castration

Next, we examined ATRA effects on MycCaP-BMP4 tumor growth in vivo. In an initial study, ATRA was found to reduce MycCaP-BMP4 tumor growth (Supplementary Fig. 1J). μCT analysis of the tumor-bearing femurs showed that ATRA led to a trend of reduced bone mineral density (BMD), percent bone volume (BV/TV) and bone surface density (BS/TV) compared with those in control non-treated mice (Supplementary Fig. 1K). However, mice treated with ATRA (12 mg/kg) led to a significant decrease in body weight at day 16 (Supplementary Fig. 1I). Dose reduction to 6 mg/kg (Supplementary Fig. 1J, arrow) alleviated the toxicity but also lessened the tumor response (Supplementary Fig. 1J, arrow). Thus, the study was repeated with several modifications (Fig. 6A). Mice were inoculated subQ+intrafemur with MycCaP-BMP4 (0.25×10^6^) and one group received ATRA (10 mg/kg) and the other received vehicle. After one week, both groups were castrated (Fig. 6A, red arrow). Castration led to a temporary halt or decrease of tumor growth in both control and ATRA-treated mice, after which tumor growth resumed in both groups and tumor growth was reduced by ATRA (Fig. 6A). μCT analysis showed that subcutaneous MycCaP-BMP4 tumors exhibited ectopic bone formation (Fig. 6B), similar to those observed with C4-2b-BMP4 tumors (Fig. 2A). ATRA reduced MycCaP-BMP4-induced bone formation measured by μCT (Fig. 6B) and Von Kossa staining (Fig. 6B).

**Figure 6.**
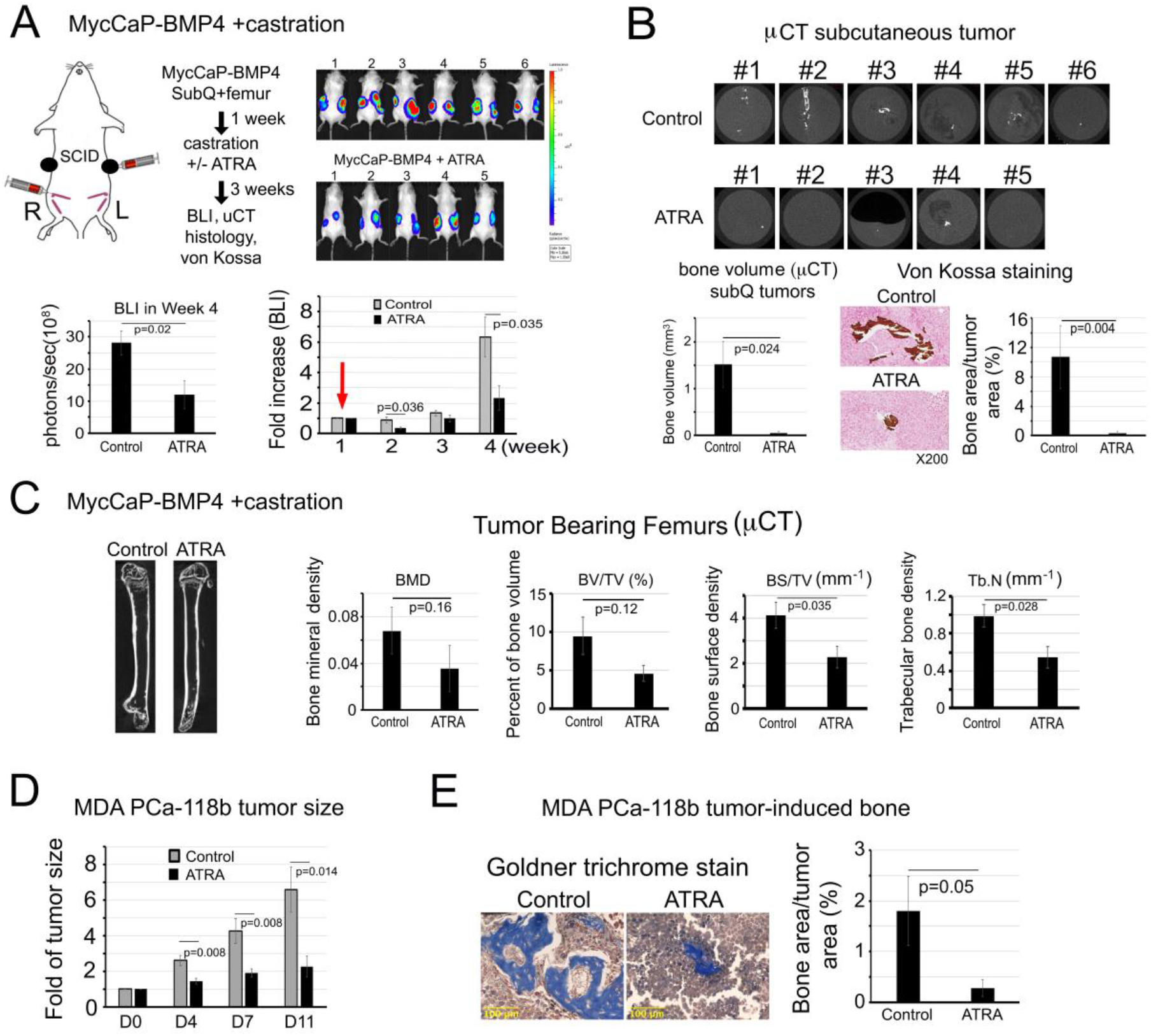
ATRA treatment reduces PCa-induced bone formation and tumor growth of MycCaP-BMP4 and MDA PCa-118b tumors. (A) MycCaP-BMP4 cells (0.25 x10^6^/site) were implanted subcutaneously and intrafemorally. Mice were castrated 1-week post-implantation (red arrow) and treated 3 weeks with or without ATRA. Tumor size measurement by BLI. Control (n=6); ATRA-treated (n=5). (B) μCT of the ectopic bone in tumors grown subcutaneously. Mineralization by von Kossa staining. (C) μCT of tumor bearing femurs from mice in (A). (D) Tumor size of MDA PCa-118b with or without ATRA treatment. Control (n=7); ATRA-treated (n=7). (E) Mineralization by Goldner Trichrome staining.

In the MycCaP-BMP4 tumor-bearing femurs, μCT analysis showed that ATRA led to a trend of decreases in bone mineral density, ratio of bone volume to total volume (BV/TV), bone surface density (BS/TV), and trabecular bone density (Tb.N) in the tumor-bearing femurs compared with those in control mice (Fig. 6C), although some parameters did not reach statistical significance. ATRA did not have an effect on MycCaP-BMP4 cell proliferation (Supplementary Fig. 1L), as also observed with Palo (Supplementary Fig. 1G). A small decrease in body weight was observed at day 28 (Supplementary Fig. 1M). Together, these results suggest that ATRA reduces MycCaP-BMP4-induced bone formation and MycCaP-BMP4 tumor growth in mice undergoing castration.

### ATRA inhibits MDA PCa-118b bone formation

MDA PCa-118b is an AR negative osteogenic PDX derived from a bone metastasis in a patient with castrate-resistant prostate cancer (40). We have previously shown that MDA PCa-118b secreted BMP4 that promotes prostate tumor growth through osteogenesis (4). We found that ATRA also reduces bone formation and tumor growth of the MDA PCa-118b xenograft (Fig. 6D).

### Palovarotene or ATRA treatments reduce plasma Tenascin C levels

In preparation for using Palo or ATRA to reduce tumor-induced bone formation through inhibition of EC-to-OSB transition in patients with castrate-resistant osteoblastic bone metastasis, we examined TNC levels, which was increased during EC-to-OSB transition, in the plasma of SCID mice with intrafemur MycCaP-BMP4 tumors treated with or without ATRA or Palo. ATRA or Palo treatments significantly reduced plasma TNC levels compared to untreated controls (Fig. 7A), suggesting that plasma TNC levels could be used to monitor response to ATRA treatment.

**Figure 7.**
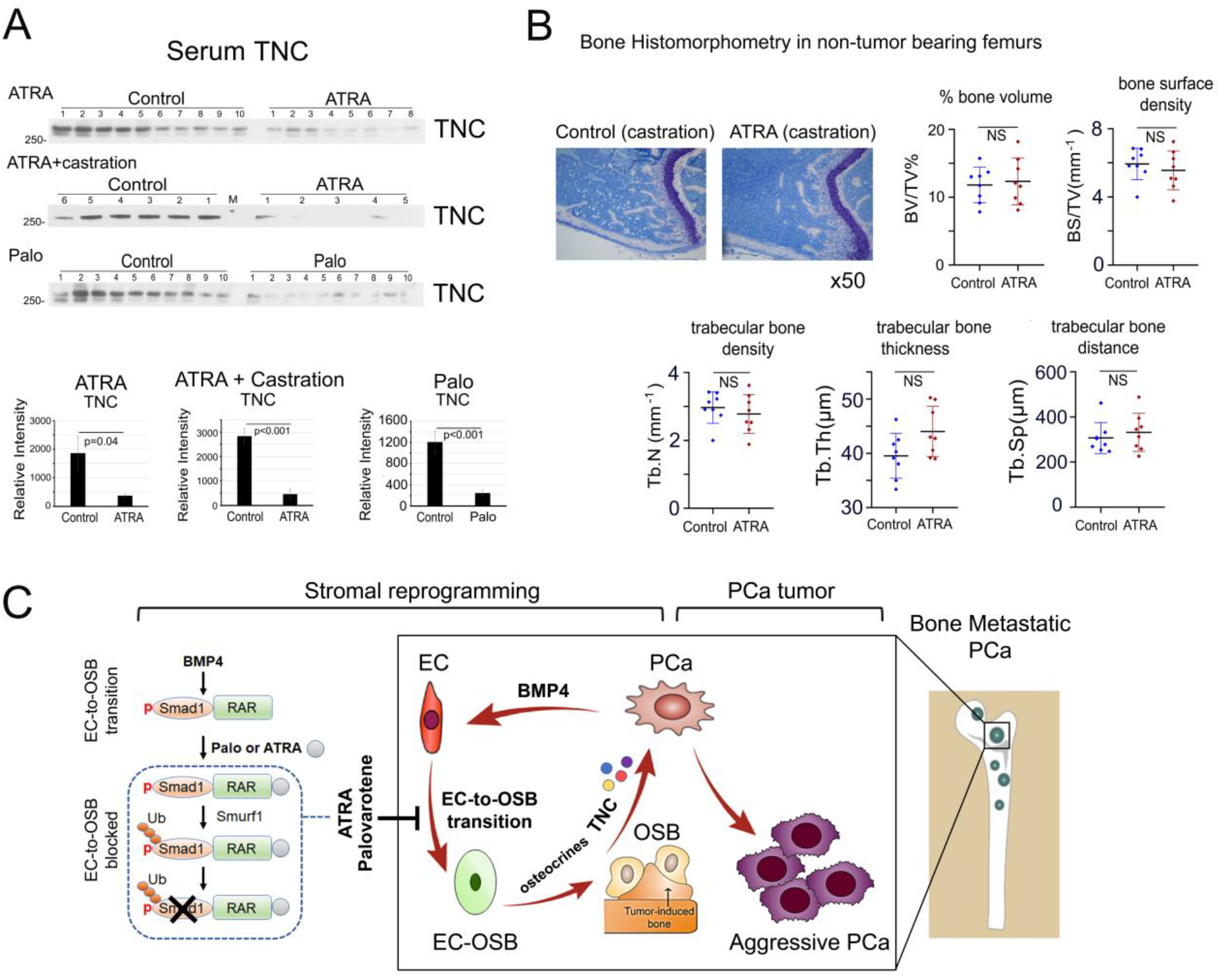
ATRA or Palo reduces plasma TNC levels in tumor-bearing mice and ATRA does not lead to overall bone loss in non-tumor involved healthy bone in castrated mice. (A) Serum TNC protein levels from SCID mice treated with ATRA only (upper), ATRA plus castration (middle), or Palo (lower) by western blot. Quantification by ImageJ. (B) Bone histomorphometry measurements in non-tumor containing femurs from ATRA plus castration-treated mice as described in Figure 6C. (C) Schematic summary. Within the tumor microenvironment, PCa-induced bone originates from ECs that have undergone EC-to-OSB transition in response to PCa-secreted BMP4 signaling through a pSmad1/RAR pathway (boxed). Activation of RARs by Palo or ATRA inhibits BMP4-mediated EC-to-OSB transition through a non-canonical RAR/pSmad1/Smurf1 pathway that results in pSmad1 degradation (dotted). This Palo/ATRA-mediated inhibitory pathway blocks stromal reprogramming and leads to a decrease in EC-OSB cell secreted factors termed “osteocrines” including TNC, a reduction in PCa-induced aberrant bone formation, and a decrease in metastatic PCa tumor growth in bone.

### ATRA does not lead to overall bone loss in non-tumor containing bones

We also examined whether ATRA leads to unwanted bone loss in healthy non-tumor-bearing bones. Bone histomorphometry analyses of contralateral non-tumor containing femurs from MycCaP-BMP4 castration plus ATRA-treated animals as described in Fig. 6C showed that there were no significant changes in bone volume (BV/TV), bone surface density (BS/TV), or trabecular bone density (Tb.N), trabecular bone distance (Tb.Sp), or trabecular thickness (Tb.Th) compared to control castrated mice (6-week-old, Fig. 7B). Similarly, no consistent changes on parameters related to activities of osteoblasts, osteoclasts, and blood vessel endothelial cells were observed (Supplementary Fig. 2). Our previous study on the effect of castration on bone showed that castration reduced bone volume and an increase in osteoclast density (41). These results suggest that ATRA treatment does not further worsen the osteoporotic effects from castration.

## Discussion

Our studies identify a strategy to target PCa-induced bone formation to interfere with the cross-talk between stromal components and metastatic PCa tumor cells. We show that activation of RAR by agonists, including Palo and ATRA, inhibits stromal reprogramming through blocking BMP4-induced EC-to-OSB transition and reduces PCa-induced aberrant bone formation and tumor growth in vivo (Fig. 7C). We also show that Palo and ATRA inhibit EC-to-OSB transition through a non-canonical RAR pathway (Fig. 7C). Because PCa-induced aberrant bone formation enhances PCa progression (4) and also resistance to therapy (6), our studies suggest that Palo or ATRA may be used to reduce tumor-induced bone formation to improve therapy outcomes for patients with PCa bone metastasis. Presently, effective therapies directed at blocking mechanisms of PCa bone metastasis are limited, with Radium-223 being the only approved agent in this category. Palo is currently being evaluated for its potential to reduce heterotopic (extra-skeletal) ossification for a rare genetic disease FOP (42). While efficacy data are pending, Palo toxicity is low (42), making its combination with other cancer-targeted therapies promising. ATRA is an FDA-approved agent for the treatment of acute promyelocytic leukemia (43). Our studies thus identify a novel therapy strategy that targets tumor-induced stromal reprogramming as treatment for PCa bone metastasis.

Our previous discovery that osteoblastic bone lesions arise from EC-to-OSB transition (3) led us to hypothesize that inhibiting this transition would represent a novel therapy strategy for bone-metastatic PCa. The current study credentials the EC-OSB transition as a valid therapy target based on ATRA or Palo-mediated inhibition of EC-to-OSB transition leading to reduced aberrant bone formation and tumor growth. This occurred without any significant effect on non-tumor involved healthy bone in castrated mice as measured by bone histomorphometry. Although ATRA and 13-cis-RA have previously been used for the treatment of primary PCa with limited or no efficacy (44–47), the potential to “repurpose” these agents in the treatment of bone metastatic PCa has not been explored. The fact that plasma levels of TNC correlated with ATRA’s effect on EC-to-OSB transition suggests their potential use as a blood-based biomarker to monitor response to ATRA treatment (Fig. 7C). Several EC-OSB cell secreted factors or “osteocrines”, including growth factors and cytokines reported in this study may also have potential as biomarkers for ATRA response. Together, our studies provide a framework for the clinical development of ATRA or Palo for the treatment of bone metastatic PCa.

TNC is an extracellular matrix glycoprotein that has been shown to initiate and sustain breast cancer metastasis to the lung (48). Our recent studies showed that TNC is highly expressed in the EC-OSB cells that rim tumor-induced bone in several osteogenic PCa bone metastasis models as well as in clinical specimens of human PCa bone metastasis (49). Further, we found that TNC increases migration, invasion and anchorage-independent growth of PCa cells in vitro and promotes PCa cell metastasis to lymph nodes and bone in vivo (49). Our finding that plasma TNC levels are reduced in response to Palo or ATRA treatment is consistent with Palo and ATRA targeting EC-to-OSB transition.

Although RARs are nuclear transcription factors, our study suggests that activation of RARs by Palo or ATRA inhibits EC-to-OSB transition via a non-canonical RAR pathway. We found that RARs form a complex with pSmad1 during BMP4-stimulated EC-to-OSB transition. It is possible that the formation of RAR/pSmad1 complex prevents the degradation of pSmad1 by Smurf1. Upon retinoic acid treatment, Smurf1 was recruited to the pSmad1/RAR/ligand complex to degrade pSmad1 (Fig. 7C). Our studies likely reveal a novel mechanism for retinoic acids’ effects on modulating skeletal development (20–22). Future studies should determine whether RARs signal through other yet-to-be identified pathways in different cellular contexts.

We found that administration of Palo or ATRA decreased the tumor volumes, raising the issues of whether this is due to a direct effect of Palo or ATRA on tumor cells or an indirect effect through inhibition of tumor-induced bone. Published data suggest that RAR agonists are not very effective in inducing growth arrest in PCa cell lines in vitro, as high concentrations (~1– 10 μM) are generally needed (50–52), instead RAR antagonists are more effective in inhibiting PCa cell proliferation in vitro (53–55). We also found that Palo or ATRA did not significantly inhibit proliferation of C4-2b-BMP4, TRAMP-BMP4, and MycCaP-BMP4 cells (Supplementary Fig. 1). When tested in vivo on subcutaneously generated non-osteogenic C4-2b and MycCaP tumors, ATRA causes only a small decrease of tumor volumes (Supplementary Fig. 3). As ATRA also inhibits angiogenesis (56), ATRA or Palo’s effects on tumor volume of osteogenic tumors are likely due to both reduction in tumor-induced bone and inhibition of tumor angiogenesis rather than a direct cytotoxic effect on tumor cells. However, the effects of ATRA or Palo on bone-forming tumor growth are moderate. Thus, we posit that ATRA or Palo in combinations with epithelial-targeting therapies (e.g. docetaxel) will be more effective for tumor growth inhibition. Our previous studies revealed that tumor-induced bone is one source of therapy resistance (6). Thus, it is also possible that Palo or ATRA, by reducing tumor-induced bone formation, will reduce resistance to therapies. Such an application requires further investigation.

In conclusion, we found that activation of RARs by agonists, including Palo and ATRA, targets the molecular basis of PCa-induced aberrant bone formation, and Palo and ATRA are promising therapeutic agents for the treatment of PCa bone metastasis.

### Materials and Methods

### Cell lines, antibodies and reagents

Cell lines, antibodies and reagents are listed in Supplementary Table 2. Cells were tested for mycoplasma using MycoAlert™ Mycoplasma Detection Kit (LONZA, LT07-418) and authenticated using short tandem repeat (STR).

### Quantitative PCR (qRT-PCR)

Total RNA was extracted using RNeasy Mini Kit (Qiagen). qRT-PCR was performed as previously described (57). Mouse-specific primer sequences are listed in Supplementary Table 3.

### Nuclear fractionation

2H11 cells were lysed in cell lysis buffer and nuclear fraction prepared as previously described (27).

### Immunoblotting, immunofluorescence, and immunoprecipitation

Cell lysates (20 μg) were analyzed by immunoblotting with indicated antibody. Western blots were quantified using ImageJ. For immunofluorescence analysis, cells were fixed and stained as previously described (58). Nuclear fractions were incubated overnight with antibodies and immunoprecipitated complexes were analyzed as described (27).

### Mineralization assays

2H11 cells were treated with BMP4 (100 ng/mL), Palo (1 μM), or both in serum-free medium for 48 h, switched into osteoblast differentiation medium, and incubated for 14 – 20 days as described (27). Cells were fixed with formalin and incubated with Alizarin Red S solution or AgNO3 solution (von Kossa staining) as described (3).

### Proximity ligation assays

Proximity ligation assay was performed using DUOLink In Situ Red Starter Kit (DUO92101) (Sigma) with anti-Rabbit PLUS (DUO92002) and anti-Mouse MINUS (DUO92004) PLA probes as previously described (58).

### RNAseq analysis

2H11 cells treated as indicated were prepared for RNAseq analysis at Arraystar Inc. (Rockville, MD). Transcriptome data have been deposited at GSE168157.

### Generation of C4-2b-BMP4-LT cell line and C4-2b-BMP4 tumors

C4-2b-BMP4-LT cell line, expressing BMP4, luciferase reporter, and red fluorescence protein Tomato, was generated as previously described (3). C4-2b-BMP4 tumors were generated in male SCID mice subcutaneously. Mice were treated by oral gavage with palovarotene (2 mg/kg/day) or corn oil (as control).

### Generation of TRAMP-BMP4-LT or MycCaP-BMP4-LC cell lines

TRAMP-BMP4 and MycCaP-BMP4 cell lines were generated by using a bicistronic retroviral vector that contains mouse BMP4 cDNA. The luciferase reporters in TRAMP-BMP4-LT and MycCaP-BMP4-LC cells were introduced using bicistronic retroviral vector pBMN-Luc-IRES-Tomato and a lentivirus vector FUW-Luc-mCh-Puro, respectively.

### Intrabone injection

TRAMP-BMP4-LT (1×10^6^) or MycCaP-BMP4-LC (0.5×10^6^) cells were injected into femurs of SCID mice. Palovarotene (2 mg/kg/day) or ATRA (10-12mg/kg/day) was administered by daily oral gavage 3 days before tumor inoculation and continued throughout the treatment period. Tumor growth was monitored weekly by bioluminescent imaging (BLI). Femurs were fixed in 10% paraformaldehyde for μCT analysis.

### Knockdown of RARα, RARβ or RARγ in 2H11 cells

RARα, RARβ, RARγ or Smurf1 in 2H11 cells were knocked down using MISSION pLKO.1 lentiviral shRNA (MilliPORE Sigma). shRNA clones were selected with 5 μg/mL puromycin. Cells infected with empty pLKO.1 lentiviral vector were used as controls. Primers for shRNA are listed in Supplementary Table 3.

### Bone histomorphometry analysis

Bone histomorphometry analysis on non-decalcified mouse femurs was performed by M.D. Anderson Cancer Center Bone Histomorphometry Core Laboratory using parameters as described (59).

### Statistical analysis

Data were expressed as the mean ± S.D. p < 0.05 by Student’s *t*-test was considered statistically significant.

### Study approval

The animal studies have been approved by the Institutional Animal Care and Use Committee at M.D. Anderson Cancer Center.

## Author contributions

G.Y., P.G.C., C.J.L., L.-Y.Y.-L., S.-H.L. conceived the idea, planned the experiments, and wrote the manuscript. G.Y., P.S., J.H.S., Y.-C.L., S.-C.L., J.P. carried out the experiments. G.Y, P.G.C., S.K.A, T.P., M.P., L.-Y.Y.-L., S.-H.L. performed data analysis and interpretation. G.Y., C.J.L., L.-Y.Y.-L., S.-H.L. provided scientific inputs for the development of the project. All the co-authors critically reviewed the present manuscript before submission.

## Acknowledgements

This work was supported by grants from the NIH (RO1 CA174798, P50 CA140388, P30 CA016672), Cancer Prevention and Research Institute of Texas (CPRIT RP150179, RP190252), and the Biology of Inflammation Center (Baylor).

## Conflict of interest

C. J. Logothetis reports receiving commercial research grants from Janssen, ORIC Pharmaceuticals, Novartis, Aragon Pharmaceuticals; and honoraria from Merck, Sharp & Dohme, Bayer, Amgen. No potential conflicts of interest are disclosed by the other authors.

---

**Supplementary Figure 1. Effect of Palo or ATRA on cell proliferation in vitro and mouse body weight in vivo in osteogenic prostate cancer models**. (A) Palo on C4-2b-BMP4 cell proliferation in vitro. (n=2). (B) Body weight of C4-2b-BMP4 tumor-bearing mice with or without Palo treatment. Control (n=5); Palo-treated (n=5). (C) BMP4 levels in TRAMP-C2 and TRAMP-BMP4 cell lines. BMP4 mRNA levels by RT-PCR or qRT-PCR. BMP4 protein levels by western blot. (D) Palo on TRAMP-BMP4-LT cell proliferation in vitro. (n=2). (E) Body weight of TRAMP-BMP4-LT tumor-bearing mice with or without Palo treatment. Control (n=7); Palo-treated (n=5). (F) BMP4 protein levels in MycCaP and MycCaP-BMP4 cell lines measured by western blot. (G) Palo on MycCaP-BMP4 cell proliferation in vitro. (n=2). (H) Body weight of MycCaP-BMP4 tumor-bearing mice with or without Palo treatment. Control (n=5); Palo-treated (n=5). (I) Body weight of MycCaP-BMP4 tumor-bearing mice treated with or without ATRA treatment. Control (n=10), ATRA-treated (n=8). (J) MycCaP-BMP4 tumor size measurement by BLI. Arrow, ATRA was reduced from 12 mg/kg to 6 mg/kg. (K) μCT analysis of tumor bearing femurs. (L) ATRA on MycCaP-BMP4 cell proliferation in vitro. (n=2). (M) Body weight of MycCaP-BMP4 tumor-bearing mice with or without ATRA plus castration. Control (n=6), ATRA-treated (n=5). *, p<0.05, **, p<0.01.

**Supplementary Figure 2. Effect of ATRA on bone histomorphometry in castrated 6-week-old mice**. Bone histomorphometry measurements in non-tumor containing femurs from ATRA plus castration-treated 6-week-old mice as described in Figure 7B. (A) Osteoblasts. (B) Endothelial cells. (C) Osteoclasts. Arrows, osteoclasts.

**Supplementary Figure 3. Effect of ATRA on the growth of subcutaneously generated non-osteogenic C4-2b, MycCaP or TRAMP-C2 tumors**. (A) MycCaP, (B) C4-2b, (C) TRAMP-C2. Tumor size was measured by BLI and expressed as fold increase over D1. Control (n=5); ATRA-treated (n=5).

**Supplementary Table 1. Secreted proteins upregulated by BMP4 but downregulated by palovarotene.**

**Supplementary Table 2. List of reagents.**

**Supplementary Table 3. Oligonucleotide sequences for qRT-PCR and shRNA knockdown.**

