## Supplemental Material for "Activation of retinoic acid receptor reduces metastatic prostate cancer bone lesions through blocking endothelial-to-osteoblast transition"

**Supplementary Material**

**Supplementary Figure 1.** Effect of Palo or ATRA on cell proliferation in vitro and mouse body weight in vivo in osteogenic prostate cancer models.

**Supplementary Figure 2.** Effect of ATRA on bone histomorphometry in castrated 6-week-old mice.

**Supplementary Figure 3.** Effect of ATRA on the growth of subcutaneously generated non-osteogenic C4-2b, MycCaP or TRAMP-C2 tumors.

**Supplementary Table 1.** Secreted proteins upregulated by BMP4 but downregulated by palovarotene.

**Supplementary Table 2.** List of reagents.

**Supplementary Table 3.** Oligonucleotide sequences for qRT-PCR and shRNA knockdown.

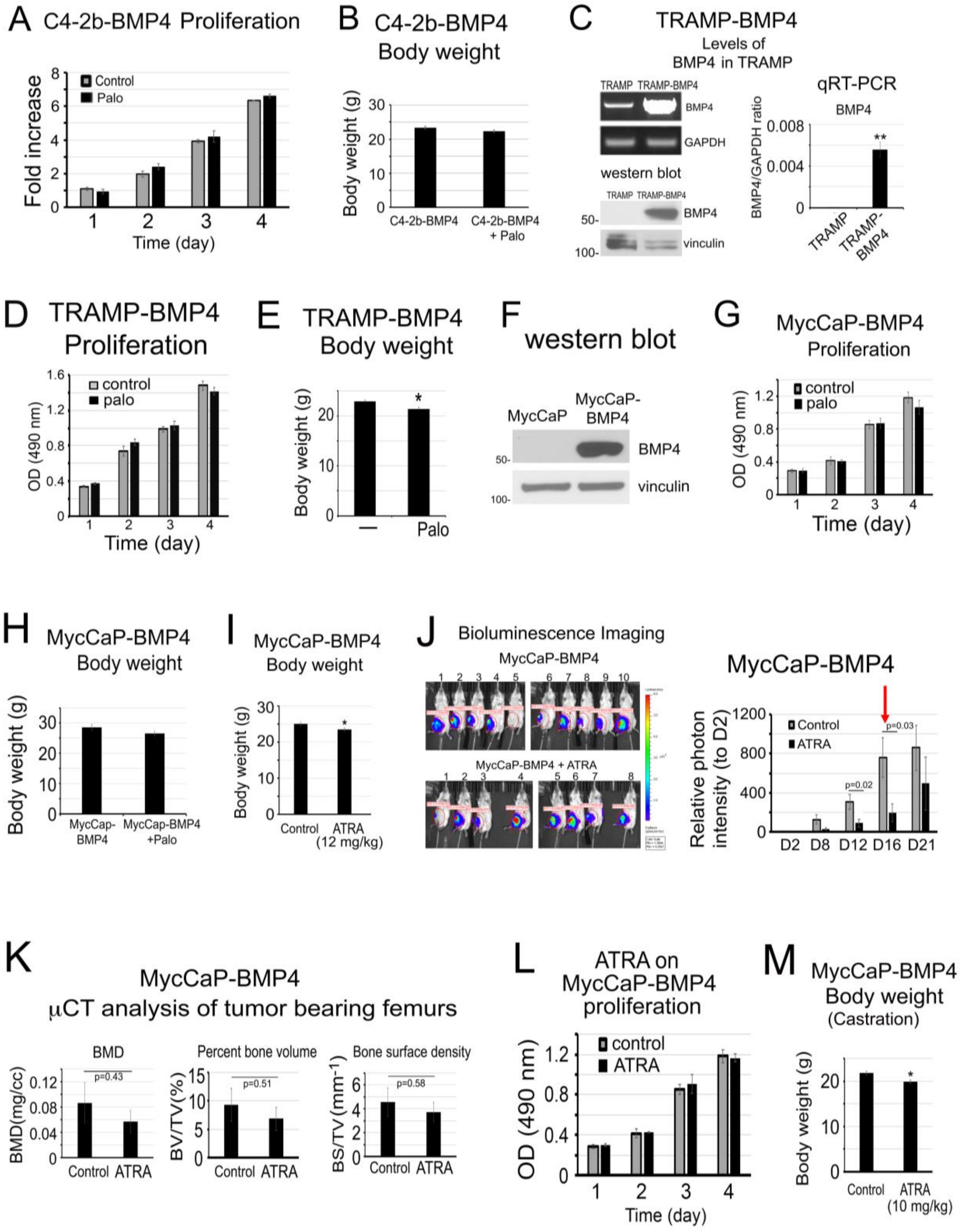


**Supplementary Figure 1. Effect of Palo or ATRA on cell proliferation in vitro and mouse body weight in vivo in osteogenic prostate cancer models.**

(A) Palo on C4-2b-BMP4 cell proliferation in vitro. (n=2). (B) Body weight of C4-2b-BMP4 tumor-bearing mice with or without Palo treatment. Control (n=5); Palo-treated (n=5). (C) BMP4 levels in TRAMP-C2 and TRAMP-BMP4 cell lines. BMP4 mRNA levels by RT-PCR or qRT-

PCR. BMP4 protein levels by western blot. (D) Palo on TRAMP-BMP4-LT cell proliferation in vitro. (n=2). (E) Body weight of TRAMP-BMP4-LT tumor-bearing mice with or without Palo treatment. Control (n=7); Palo-treated (n=5). (F) BMP4 protein levels in MycCaP and MycCaP-BMP4 cell lines measured by western blot. (G) Palo on MycCaP-BMP4 cell proliferation in vitro. (n=2). (H) Body weight of MycCaP-BMP4 tumor-bearing mice with or without Palo treatment. Control (n=5); Palo-treated (n=5). (I) Body weight of MycCaP-BMP4 tumor-bearing mice treated with or without ATRA treatment. Control (n=10), ATRA-treated (n=8). (J) MycCaP-BMP4 tumor size measurement by BLI. Arrow, ATRA was reduced from 12 mg/kg to 6 mg/kg. (K) µCT analysis of tumor bearing femurs. (L) ATRA on MycCaP-BMP4 cell proliferation in vitro. (n=2). (M) Body weight of MycCaP-BMP4 tumor-bearing mice with or without ATRA plus castration. Control (n=6), ATRA-treated (n=5). *, $p<0.05$, **, $p<0.01$.

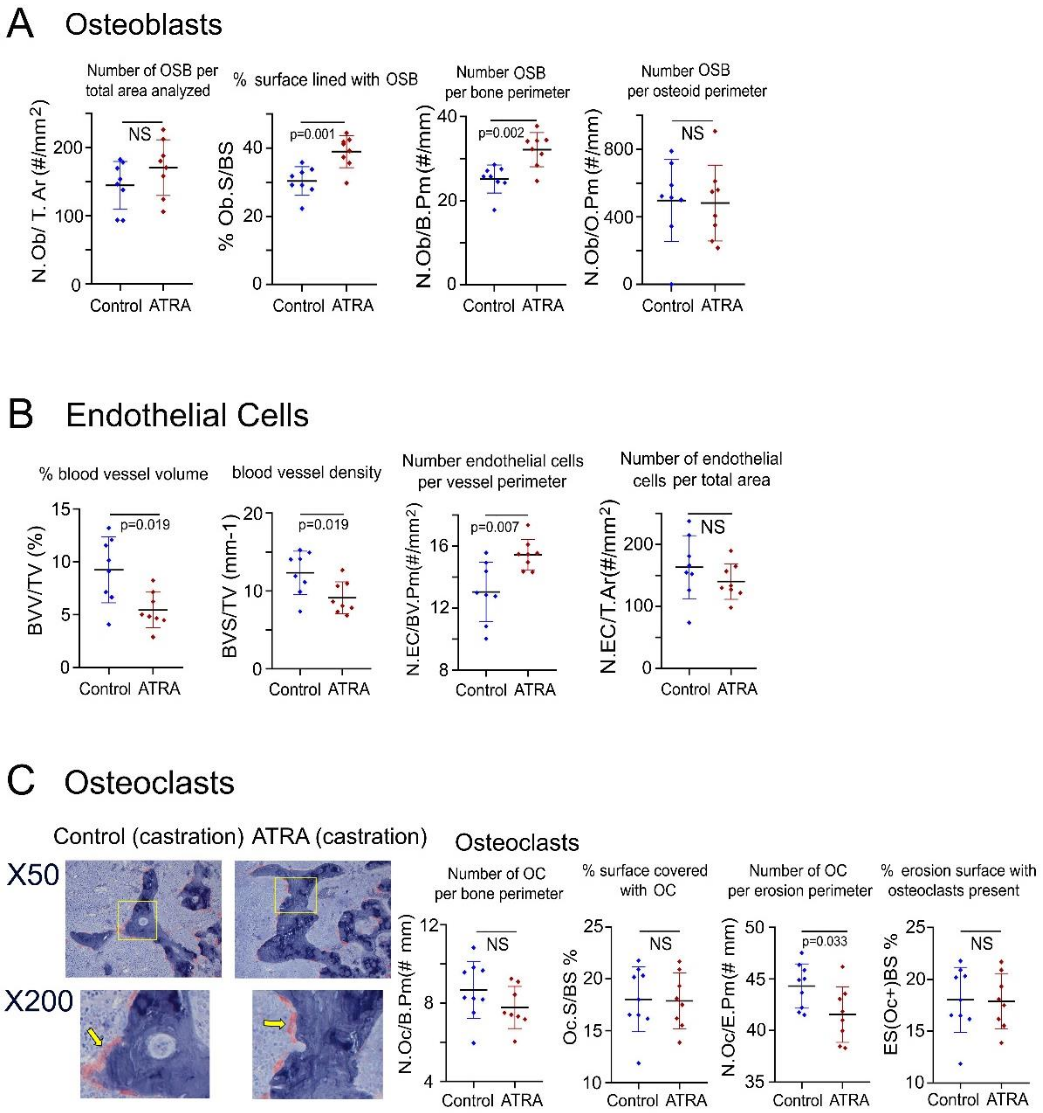


**Supplementary Figure 2. Effect of ATRA on bone histomorphometry in castrated 6-week-old mice.**

Bone histomorphometry measurements in non-tumor containing femurs from ATRA plus castration-treated 6-week-old mice as described in Figure 7B. (A) Osteoblasts. (B) Endothelial cells. (C) Osteoclasts. Arrows, osteoclasts.

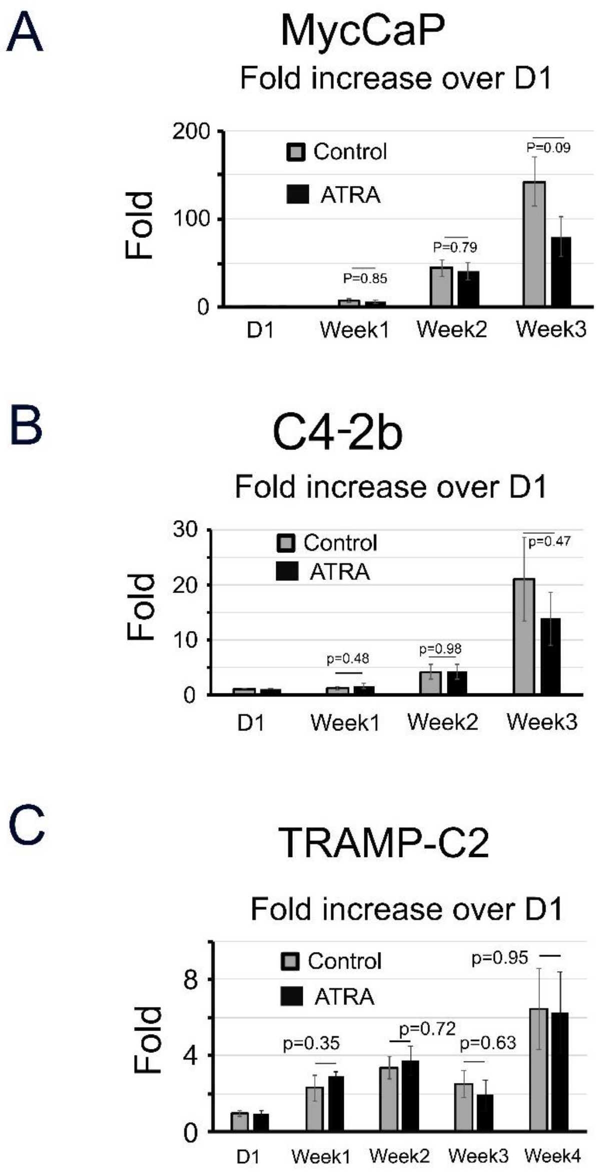


**Supplementary Figure 3. Effect of ATRA on the growth of subcutaneously generated non-osteogenic C4-2b, MycCaP or TRAMP-C2 tumors.**

(A) MycCaP, (B) C4-2b, (C) TRAMP-C2. Tumor size was measured by BLI and expressed as fold increase over D1. Control (n=5); ATRA-treated (n=5).

**Supplementary Table 1. Secreted proteins upregulated by BMP4 but downregulated by palovarotene.**

| Symbol | Entrez Gene Name | Palo/BMP4 (Fold) |
|---|---|---|
| **Growth factor** | | |
| FGF10 | fibroblast growth factor 10 (Fgf10) | 0.222707424 |
| HGF | hepatocyte growth factor (Hgf) | 0.445753255 |
| Prl2c2 | prolactin family 2, subfamily c, member 2 (Prl2c2) | 0.478084416 |
| FGF11 | fibroblast growth factor 11(Fgf11) | 0.609049314 |
| **Cytokines** | | |
| TSLP | thymic stromal lymphopoietin (Tslp) | 0.267576076 |
| WNT4 | Wnt family member 4 (Wnt4) | 0.464882943 |
| WNT5A | Wnt family member 5A (Wnt5a) | 0.484665661 |
| CXCL14 | C-X-C motif chemokine ligand 14 (Cxcl14) | 0.534851622 |
| **Peptidase** | | |
| Masp1 | mannan-binding lectin serine peptidase 1 (Masp1) | 0.271255061 |
| ADAMTS9 | ADAM metallopeptidase with thrombospondin type 1 motif 9(Adamts9) | 0.456168831 |
| PRSS12 | serine protease 12 (Press12) | 0.576781327 |
| **Enzyme** | | |
| LOX | lysyl oxidase (Lox) | 0.349213911 |
| FN1 | fibronectin 1 (Fn1) | 0.531306991 |
| **Others** | | |
| MGP | matrix Gla protein (Mgp) | 0.087132725 |
| GREM1 | gremlin 1, DAN family BMP antagonist (Grem1) | 0.108909574 |
| BGLAP | bone gamma-carboxyglutamate protein (Bglap2) | 0.136125654 |
| FMOD | fibromodulin (Fmod) | 0.169962587 |
| GREM2 | gremlin 2, DAN family BMP antagonist (Grem2) | 0.170140442 |
| PCOLCE2 | procollagen C-endopeptidase enhancer 2 (Pcolce2) | 0.200583658 |
| THBS1 | thrombospondin 1 (Thbs1) | 0.202858139 |
| CCK | cholecystokinin (Cck) | 0.217947602 |
| Cdsn | corneodesmosin (Cdsn) | 0.280749707 |
| VCAN | versican (Vcan) | 0.287769784 |
| SERPINE2 | serpin family E member 2 (Serpine2) | 0.288900152 |
| COL12A1 | collagen type XII alpha 1 chain (Col12a1) | 0.333460366 |
| PTX3 | pentraxin 3 (Ptx3) | 0.444080026 |
| C1QTNF3 | C1q and TNF related 3 (C1qtnf3) | 0.444715731 |
| TNC | tenascin C (Tnc) | 0.461806831 |
| KAZALD1 | Kazal type serine peptidase inhibitor domain 1 (Kazald1) | 0.469319271 |
| PRELP | proline and arginine rich end leucine rich repeat protein (Prelp) | 0.500851789 |
| NSMF | NMDA receptor synaptonuclear signaling and neuronal migration factor (Nsmf) | 0.508656716 |
| BGN | biglycan (Bgn) | 0.54390398 |
| PODNL1 | podocan like 1 (Podnl1) | 0.561904762 |
| IGFBP2 | insulin like growth factor binding protein 2 (lgfbp2) | 0.566866267 |
| SASH1 | SAM and SH3 domain containing 1 (Sash1) | 0.613924051 |
| COL10A1 | collagen type X alpha 1 chain (Col10a1) | 0.616504854 |
| APLN | apelin (Apln) | 0.625854568 |
| BBS9 | Bardet-Biedl syndrome 9 (Bbs9) | 0.657033806 |
| THSD1 | thrombospondin type 1 domain containing 1 (Thsd1) | 0.660432496 |

**Supplementary Table 2. List of reagents.**

| Reagent | Source | Catalogue Number |
|---|---|---|
| **Antibodies** | | |
| BMP4 | Santa Cruz Biotechnology | SC-12721, RRID: AB_2063534 |
| OSX | AVIVA SYSTEMS BIOLOGY | ARP32231_P050; RRID: AB_10638646 |
| Tenascin C | MilliPORE Sigma | AB19011; RRID: AB_2203804 |
| Smurf1 | Santa Cruz Biotechnology | sc-100616; RRID:AB_1129557 |
| Smurf1 | Cell Signaling Technology | # 2174; RRID:AB_2193845 |
| RARα | Cell Signaling Technology | # 62294, RRID:AB_2799625 |
| RARβ | Thermo Fisher Scientific | MA1-811, RRID:AB_ |
| RARγ | Cell Signaling Technology | # 8965, RRID:AB_10998934 |
| Phospho-Smad1/5(Ser463/465) | Cell Signaling Technology | # 9516, RRID:AB_491015 |
| Smad1(D59D7) | Cell Signaling Technology | #6944, RRID:AB_10858882 |
| GAPDH(D16H11) | Cell Signaling Technology | # 5174, RRID:AB_10622025 |
| Vinculin(E1E9V) | Cell Signaling Technology | # 13901, RRID:AB_2728768 |
| LaminA/C | Cell Signaling Technology | #2032S; RRID:AB_2136278 |
| Histone H3 | Cell Signaling Technology | #4499S; RRID:AB_10544537 |
| **Inhibitors** | | |
| Palovarotene | Toronto Research Chemicals | P165900 |
| ATRA | Millipore-Sigma | S1574 |
| 13-cis RA | Selleckchem | CT99021 |
| **Recombinant Protein** | | |
| Recombinant human BMP4 | R&D Systems | 314-BP |
| Mouse TNC ELISA Kit | MyBioSource | #MBS9627232 |
| Mouse Osteocalcin Enzyme Immunoassay Kit | Alfa Aesar | BT-470 |
| **Cell Lines** | | |
| 2H11 | ATCC Cat# CRL-2163 | RRID:CVCL_6762 |
| MycCaP | ATCC Cat# CRL-3255 | RRID:CVCL_J703 |
| TRAMP-C2 | ATCC Cat# CRL-2731 | RRID:CVCL_3615 |

**Supplementary Table 3. Oligonucleotide sequences for qRT-PCR and shRNA knockdown.**

| Name | Sequence (5'-3') | product length(bp) |
|---|---|---|
| OSX-qPCR-F | CTTCCCTGGATATGACTCAT | |
| OSX-qPCR-R | TGTCCCACCAAGGAGTAGGT | 165 |
| Osteocalcin-qPCR-F | GCTCTGTCTCTCTGACCTCA | |
| Osteocalcin-qPCR-R | TGGACATGAAGGCTTTGTCA | 68 |
| RARα-qPCR-F | GGGCGAGATGTACGAGAGTG | |
| RARα-qPCR-R | TCGATGGAGTGGTTTGAGCC | 185 |
| RARβ-qPCR-F | CTCAGATGCACAATGCTGGC | |
| RARβ-qPCR-R | TGCTTCCAGCAGTGGTTCTT | 191 |
| RARγ-qPCR-F | CTCAGCACTGCCTTTCGGAT | |
| RARγ-qPCR-R | GAGGTACAGACGGGCTCACATT | 325 |
| Slug-qPCR-F | CTGTATGGACATCGTCGGCA | |
| Slug-qPCR-R | ACTTACACGCCCCAAGGATG | 95 |
| Foxo1-qPCR-F | TCAAGGATAAGGGCGACAGC | |
| Foxo1-qPCR-R | CCATGGACGCAGCTCTTCTC | 183 |
| Maf-qPCR-F | TGTTAAATGCTCCGTGGGGG | |
| Maf-qPCR-R | CCCGGTTCAAAGGTGAGCTA | 419 |
| Cited2-qPCR-F | AAAAGGGAACGGCTCGGAAT | |
| Cited2-qPCR-R | GCGATTTCTGCTCGGAACAC | 160 |
| Foxf2-qPCR-F | TACCATCGCGTGGTGAGCG | |
| Foxf2-qPCR-R | GTGGAGTGGTGCTGGTAACG | 590 |
| Fmn1-qPCR-F | AGTTGCCACCAGAATACCAGG | |
| Fmn1-qPCR-R | CGGCGCTTTCCTAACTGTTG | 181 |
| Id4-qPCR-F | AAACAAGCCACCGGAGGAAA | |
| Id4-qPCR-R | CGTACGGTGAATGCTCGTGA | 172 |
| Tnc-qPCR-F | CTCTGGAATTGCTCCCAGCAT | |
| Tnc-qPCR-R | TTCCGGTTCAGCTTCTGTGGTAG | 70 |
| Fgf10-qPCR-F | CCGACACCACCAGTTCCTAC | |
| Fgf10-qPCR-R | CTTTGACGGCAACAACTCCG | 656 |
| Tslp-qPCR-F | TTTGCCCGGAGAACAAGAGA | |
| Tslp-qPCR-R | TAGAGTAAGGTGTGTGCAGGG | 428 |
| Mgp-qPCR-F | CAGACTCACAGGACACCCGA | |
| Mgp-qPCR-R | GCTTTAGCTCGCCACCTCTG | 187 |
| RunX2-qPCR-F | TGCTCACTCCGTTTTGTTTTGTT | |
| RunX2-qPCR-R | CCCATCTGGTACCTCTCCGA | 687 |
| GAPDH-qPCR-F | TGCAGTGGCAAAGTGGAGAT | |
| GAPDH-qPCR-R | TTTGCCGTGAGTGGAGTATA | 96 |
| mBMP4-BamH1-F | GGATCCATGATTCCTGGTAACCGAATGCTG | |
| mBMP4-EcoR-H-R | GAATTCTCATCAGCGGCATCCACACCCCTCT | 1224 |

### shRNA

| Gene | Accessions | Clone ID | Mature Antisense (5'-3') | Target |
|---|---|---|---|---|
| RARα | NM_009024 | TRCN0000054591(#1) | CCGGGCTGGAAGCACTGAAAGTCTACTCGAGTAGACTTTCAGTGCTTCCAGCTTTTTG | CDS |
| | NM_009024 | TRCN0000027125(#4) | CCGGCTGAAAGTCTACGTCCGGAAACTCGAGTTTCCGGACGTAGACTTTCAGTTTTT | CDS |
| RARβ | NM_011243 | TRCN0000027071 (#3) | CCGGCCAAGTTCAAGTGGGAATATACTCGAGTATATTCCCACTTGAACTTGGTTTTT | CDS |
| | NM011243 | TRCN0000027121 (#5) | CCGGGCATGTCCAAAGAGTCTGTTACTCGAGTAACAGACTCTTTGGACATGCTTTTT | CDS |
| RARγ | NM_011244 | TRCN0000027093 (#1) | CCGGCAATGACAAGTCTTCTGGCTACTCGAGTAGCCAGAAGACTTGTCATTGTTTTT | CDS |
| | NM_011244 | TRCN0000279129 (#3) | CCGGTGCGGATCTGTACAAGGTATACTCGAGTATACCTTGTACAGATCCGCATTTTTG | CDS |
| Smurf1 | NM_029438 | TRCN0000040573 (#1) | CCGGCCAGTATTCCACGGACAATATCTCGAGATATTGTCCGTGGAATACTGGTTTTTG | CDS |
| | NM_029438 | TRCN0000040573 (#3) | CCGGGCTGGATAAGATAGACCTGAACTCGAGTTCAGGTCTATCTTATCCAGCTTTTTG | CDS |
